# The safety profile of Bald’s eyesalve for the treatment of bacterial infections

**DOI:** 10.1101/2020.04.23.041749

**Authors:** Blessing O Anonye, Valentine Nweke, Jessica Furner-Pardoe, Rebecca Gabrilska, Afshan Rafiq, Faith Ukachukwu, Julie Bruce, Christina Lee, Meera Unnikrishnan, Kendra P. Rumbaugh, Lori AS Snyder, Freya Harrison

## Abstract

The rise in antimicrobial resistance has prompted the development of alternatives, such as plant-derived compounds, to combat bacterial infections. Bald’s eyesalve, a remedy used in the Early Medieval period, has previously been shown to have efficacy against *Staphylococcus aureus* grown in an *in vitro* model of soft tissue infection. This remedy also had bactericidal activity against methicillin-resistant *S. aureus* (MRSA) in a chronic mouse wound. However, the safety profile of Bald’s eyesalve has not yet been demonstrated, and this is vital before testing in humans. Here, we determined the safety potential of Bald’s eyesalve using *in vitro*, *ex vivo*, and *in vivo* models representative of skin or eye infections. We also confirmed that Bald’s eyesalve is active against an important eye pathogen, *Neisseria gonorrhoeae*. Low levels of cytotoxicity were observed in eyesalve-treated cell lines representative of skin and immune cells. Results from a bovine corneal opacity and permeability test demonstrated slight irritation to the cornea that resolved within 10 minutes. The slug mucosal irritation assay revealed that a low level of mucus was secreted by slugs exposed to eyesalve, indicating mild mucosal irritation. We obtained promising results from mouse wound closure experiments; no visible signs of irritation or inflammation were observed. Our results suggest that Bald’s eyesalve could be tested further on human volunteers to assess safety for topical application against bacterial infections.

**Importance:** Alternative treatment for bacterial infections are needed to combat the ever increasing repertoire of bacteria resistant to antibiotics. A medieval plant-based remedy, Bald’s eyesalve, shows promise as a substitute for the treatment of these infections. For any substance to be effective in the treatment of bacterial infections in humans, it is important to consider the safety profile. This is a key consideration in order to have the necessary regulatory approval. We demonstrate the safety profile of Bald’s eyesalve using a variety of models, including whole-organ and whole-animal models. Our results show that Bald’s eyesalve is mildly toxic to cultured human cells, but potentially suitable for patch tests on healthy human volunteers to assess safety for later clinical trials. Our work has the potential to transform the management of diseases caused by bacterial infections, such as diabetic foot ulcers, through topical application of a natural product cocktail based on Bald’s eyesalve.

## Introduction

There is an urgent need for the development of new antimicrobials to fight bacterial infections which have become resistant to antibiotics. Antimicrobial resistance (AMR) now poses a serious threat to human health, especially to those with a suppressed immune system. AMR has an associated mortality of over 33,000 people per annum in Europe, with a prediction that 10 million people per year are likely to be killed by 2050 globally (1–4). This rise in AMR, coupled with the lack of development of new antibiotics, has prompted research into the use of alternatives, such as plant-derived compounds (5–7).

Historically, plant-based compounds have been explored for their antimicrobial properties. These include oils from *Cupressus sempervirens* (Cypress) and *Commiphora* species (myrrh), being used to treat colds, coughs, and inflammation (8), and *Artemisia annua* (Artemisin) used to treat malaria (9). Bald’s eyesalve has previously been shown to possess antibacterial activity against *Staphylococcus aureus* in planktonic cultures and biofilms (10), and against *Pseudomonas aeruginosa* in planktonic cultures (11). A recent study from our laboratory found that Bald’s eyesalve had antimicrobial activity against a range of Gram-positive and Gram-negative organisms in planktonic cultures and biofilms (12). However, the safety profile of the eyesalve has not previously been reported. To explore whether this eyesalve could potentially be effective against bacterial infections in humans, it is important to firstly determine the safety profile.

Traditionally, animals have been employed for safety testing of compounds, but in order to reduce the number of animals used in line with the 3Rs (replacement, reduction and refinement of animal use) (13), we decided to use alternate models. Cell lines are particularly robust for measuring the cytotoxic effects of drug formulation and appropriate cell lines been used as surrogates for different body sites. Human cell lines, HaCaT and THP-1, have been used as substitutes for human skin cells and immune cells respectively. These cell lines have been employed for cytotoxicity testing of other natural products, drugs, and cosmetic products (13–16).

The bovine corneal opacity and permeability (BCOP) test is an eye irritation assay that replaced the Draize rabbit test (an *in vivo* assay) which identifies chemicals/irritants that induce serious eye damage (17, 18). It is described in Section 4 of the Organisation for Economic Co-operation and Development’s guidelines (Test No 437, https://doi.org/10.1787/9789264203846-en) for assays that can be used to determine the health effects of chemicals. The BCOP assay assesses the effect of potential irritants on the opacity and permeability of the isolated corneas. Toxicity of the test substance is measured by decreased light transmission leading to opacity, indicating corneal injury, and increased fluorescein dye permeability, reflecting damage to the corneal epithelium. This assay acts as an alternative to the use of animals in initial toxicity testing as the isolated bovine eyes are abattoir by-products that would otherwise be disposed of. Churchward and colleagues used the BCOP test to determine that antimicrobial fatty acids proposed for use in prophylaxis and treatment of gonococcal eye infections do not cause irritation (19). It was also employed to test tacrolimus, an immunosuppressive drug for the treatment of autoimmune-based inflammatory eye disorders (20).

Similarly, the slug mucosal irritation (SMI) assay is an alternative to the Draize test that has been used in testing the eye irritation potency of ingredients in cosmetic products, such as shampoos (21). The premise of the SMI test is that mucus will be produced when slugs are exposed to irritating substances and the slugs will lose weight. Hence, irritants will cause weight loss, mucus production, and release of proteins and enzymes due to tissue damage (22). This can be indirectly correlated to the effects of substances on humans, such as the stinging, itching and burning normally experienced when eyes are irritated, leading to red and watery eyes (21).

The aim of this present study was to determine the safety and irritation potential of Bald’s eyesalve using alternative *in vitro* and *ex vivo* models to consider safety parameters before undertaking testing on humans. Human cell lines were employed as an initial screen, followed by the BCOP test and slug mucosal irritation assays. Given the original prescription of the remedy was to treat eye infections, we also confirmed the activity of the eyesalve against *Neisseria gonorrhoeae*, a common cause of neonatal conjunctivitis. Finally, due to the low levels of cytotoxicity observed in our *in vitro* and *ex vivo* assays, we performed wound healing experiments in mouse models. This was necessary as our research is primarily geared towards the topical application of the eyesalve, or application of the active ingredients therein, in the treatment of bacterial infections such as those affecting the skin and eyes.

## Results

### Bald’s eyesalve is active against *Neisseria gonorrhoeae*

We have previously shown that Bald’s eyesalve is active against *S. aureus* in planktonic culture, in biofilms formed in an artificial model of wound infection, and in biopsies taken from infected mouse wounds (10). Here, we wanted to investigate a further eye pathogen of clinical interest. We choose *N. gonorrhoeae* as it is a pathogen that causes neonatal conjunctivitis. We performed disk diffusion assays using sterile 6 mm disks infused with 10 μl of three batches of eyesalve placed on a gonococcal agar plate containing the multi-drug resistant *N. gonorrhoeae* NCCP11945 strain (Chung *et al.*, 2008). The results showed that Bald’s eyesalve was active against this strain as significantly larger zones of inhibition were obtained compared to the control (Figure 1A, ANOVA, F_3,12_ = 13729, p < 0.001). Furthermore, planktonic cultures of this strain were exposed briefly to the three batches of the eyesalve and a 7-log reduction of *N. gonorrhoeae* was observed compared to the control, as no viable colonies were found on the eyesalve-treated cultures (Figure 1B).

**Figure 1:**
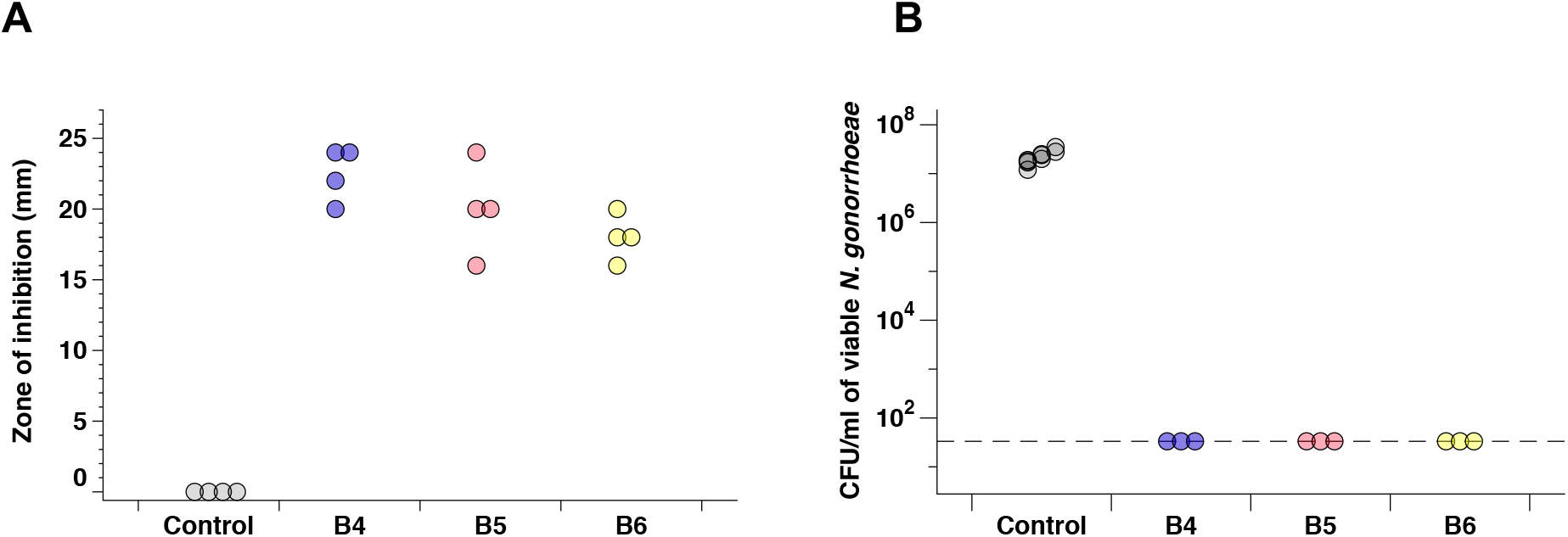
Antibacterial activity of Bald’s eyesalve against *N. gonorrhoeae*. A) Zones of inhibition of three batches of Bald’s eyesalve (B4, B5 and B6) against *N. gonorrhoeae* NCCP11945, control is sterile distilled water. We found differences between the control and the three eyesalve batches, ANOVA F_3,12_ = 13729,, p < 0.001. B) The number of colonies obtained after treatment with three batches of eyesalve, control is PBS. The dashed line represents the limit of detection. We found a difference between the control and the three eyesalve batches, ANOVA, F_3,14_ = 267.66, p < 0.001.

### The viability of human cells treated with Bald’s eyesalve

The lack of toxicity of any natural product is a key requirement for applications in humans, so we first determined the toxicity when applied to human cells. Two cell lines, HaCaT and THP-1, were treated with Bald’s eyesalve, and alamarBlue™ was used to assess the viability of the cells. Viable cells convert the oxidised form of alamarBlue™ into the reduced form with a change in colour from blue to pink, which is then measured to determine the percentage of cells which are viable (% alamarBlue reduced). As a control, we also treated cells with Neosporin^®^ or Optrex™ chloramphenicol eye drops. Both of these preparations are available without prescription in the USA and the UK. While safe for topical use, chloramphenicol is known to cause death by apoptosis when applied to cell cultures (23). For the HaCaT cells, treatment with the undiluted eyesalve led to the death of the cells with a similar result in the chloramphenicol control treated cells (Figure 2A, 2B). More viable HaCaT cells were observed in the 1/10 dilution of the eyesalve compared to the undiluted eyesalve. However, the diluted chloramphenicol control had significantly higher viable cells than the diluted eyesalve (ANOVA, F^9,21^ = 6293, p < 0.04, Figure 2A). Whereas, the viability of THP-1 cells treated with both the undiluted and diluted eyesalve were similar (Figure 2C, 2D), and significantly more viable cells also observed in the chloramphenicol treated controls (ANOVA, F_9,30_ = 66.77, p < 0.001, Figure 2D).

**Figure 2:**
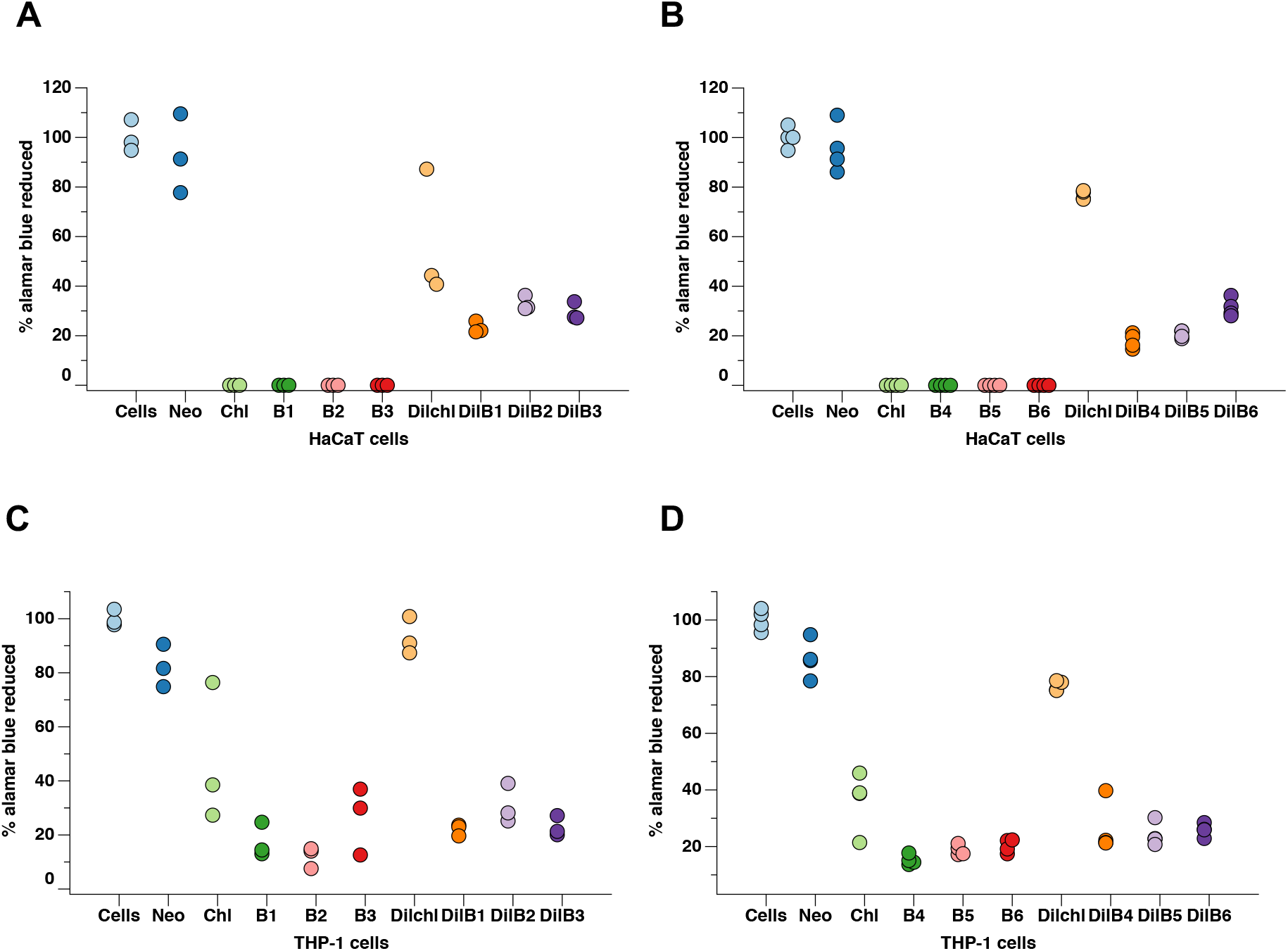
AlamarBlue™ assay to determine the viability of cells treated with eyesalve. A) HaCaT cells were treated with three batches of undiluted and 1/10 dilution of the eyesalve (B1, B2, B3) for 24 hours and B) treated with another three batches, B4, B5 and B6 for 24 hours. The controls include cells only (untreated), Neosporin (Neo), a safe antibiotic for wound infections and Optrex™ chloramphenicol (chl) treated cells (n = 3-4 replicates). The preface “dil” represents cells treated with a 1/10 dilution of either the chloramphenicol or the different eyesalve batches. ANOVA found a significant difference between the diluted chloramphenicol and diluted eyesalve batches followed by Dunnett’s test for multiple comparison, F_8,27_ = 25432, p < 0.001. C) THP-1 cells were treated with three batches of undiluted and 1/10 dilution of the eyesalve (B1, B2, B3) for 24 hours. No significant difference observed between the diluted and undiluted eyesalve (ANOVA, F_9,20_ = 18.85, p > 0.1) D) THP-1 cells treated with three separate batches, B4, B5 and B6 for 24 hours. A significant difference was found between the chloramphenicol treated cells and eyesalve treated cells, followed by Dunnett’s test for multiple comparison (ANOVA, F_9,30_ = 66.77, p < 0.001).

Next, we measured the levels of lactate dehydrogenase (LDH) released by the cells after treatment with Bald’s eyesalve. LDH is a cytoplasmic enzyme found in cells and is released when the plasma membrane is damaged due to cells undergoing necrosis, apoptosis or other cellular damage (24). Treatment of HaCaT cells with the undiluted eyesalve for 24 hours led to low levels of LDH being released but this was not significantly different from the cell only control (ANOVA, F_9,30_ = 18.19, p > 0.9, Figure 3A). However, treatment of the HaCaT cells with 1/10 chloramphenicol released more LDH compared to the eyesalve indicating more damage to the plasma membrane (ANOVA, F_9,30_ = 18.19, p < 0.001, Figure 3A, 3B). We initially treated the THP-1 cells with three batches of eyesalve and did not observe any LDH being released, including the chloramphenicol treated cells (Figure S1). We then tested an additional three eyesalve batches in both the undiluted and diluted forms (Figure 3C). Similar to the HaCaT cells, THP-1 cells treated with a 1/10 dilution of the chloramphenicol led to a significantly higher level of LDH compared to the 1/10 eyesalve treated cells (ANOVA, F_9,40_ = 1817, p < 0.001). These results further demonstrate that while the eyesalve clearly causes some damage to cultured human cells, it may be less cytotoxic than the chloramphenicol eye drops used for the treatment of conjunctivitis.

**Figure 3:**
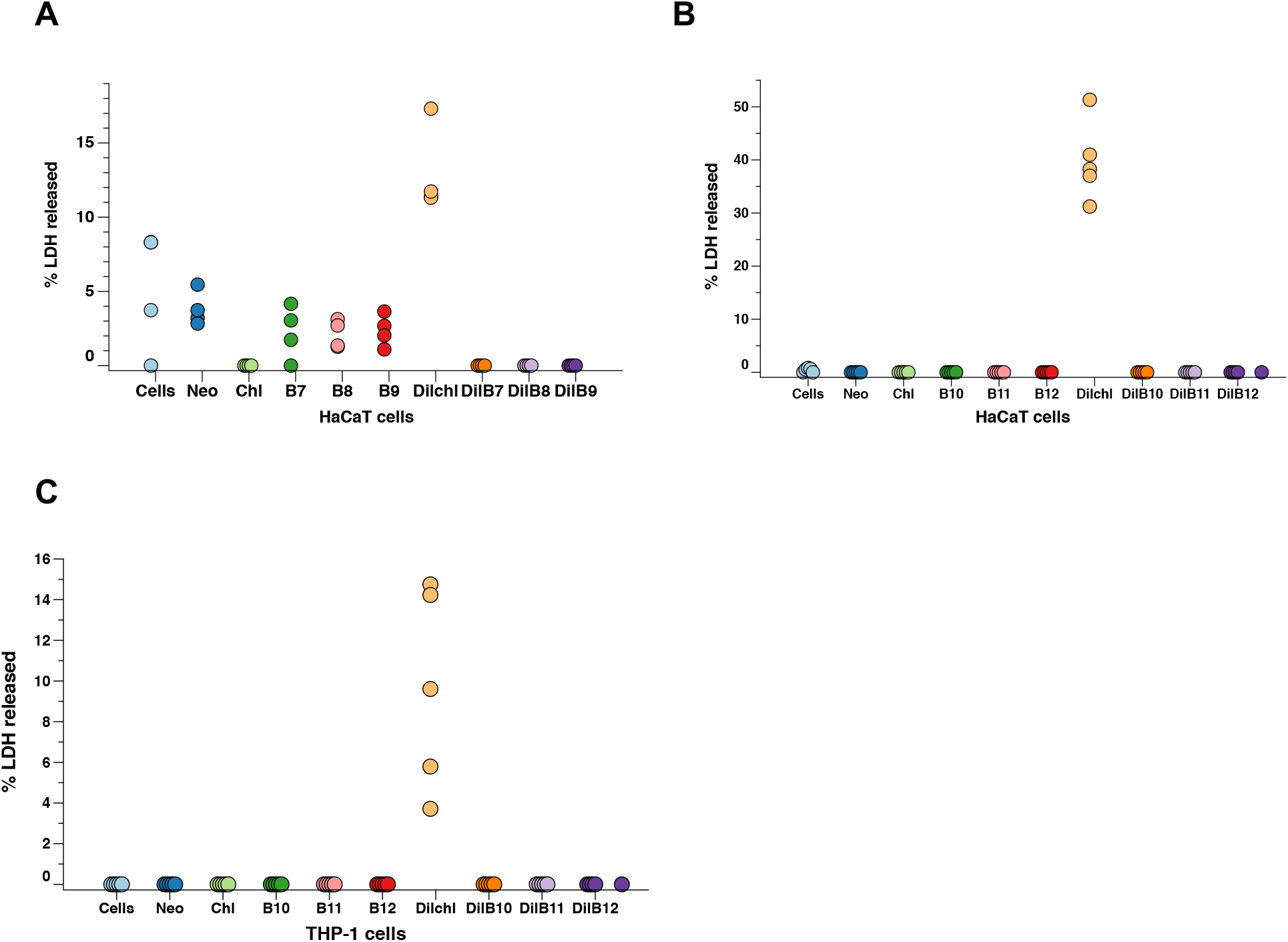
Lactate dehydrogenase assay to determine the cytotoxicity profile of the eyesalve. A, B) HaCaT cells were treated with six batches of the eyesalve. A significant difference was observed in cells treated with 1/10 dilution of the eyesalve compared to the 1/10 chloramphenicol treated cells by ANOVA. This was followed by Dunnett’s test for multiple comparison, F_9,30_ = 18.19, p < 0.001, n = 4-5 replicates. C) THP-1 cells were treated with three batches of diluted eyesalve (B10, B11 and B12) for 24 hours and the extracellular LDH released was measured. ANOVA found a significant difference in cells treated with 1/10 dilution of the eyesalve compared to the 1/10 chloramphenicol treated cells followed by Dunnett’s test for multiple comparison (F_9,40_ = 1817, p < 0.001, n = 5 replicates) The controls include cells only (untreated), Neosporin (Neo), a safe antibiotic for wound infections and Optrex™ chloramphenicol (chl) treated cells (n = 4-5 replicates). The preface “dil” represents cells treated with a 1 in 10 dilution of either the chloramphenicol or the different eyesalve batches.

### Bald’s eyesalve causes a brief irritation in the bovine corneal opacity & permeability assay

Bovine eyes were spotted with the different batches of eyesalve and observed for opacity of the corneal surface followed by washing with PBS. Each eye was then stained with fluorescein and visualized under a cobalt blue filter to reveal any damage and increased permeability caused by the eyesalve. Treatment of the bovine eyes with six batches of eyesalve revealed a transient change to the surface of the eye, which may indicate irritation of the cornea that falls below the criteria for BCOP scoring. The observed change, a slight opacity, resolved within the 10 minutes incubation of the bovine eye prior to fluorescein staining and was not present at the scoring stage, although opacity and fluorescein staining could be seen in the positive control, (0.5M NaOH) a strong irritant (Figure 4). Examination of the corneas for opacity and epithelial damage at the conclusion of the BCOP protocol demonstrated that the six batches of eyesalve caused no irritation (Table 1).

**Figure 4:**
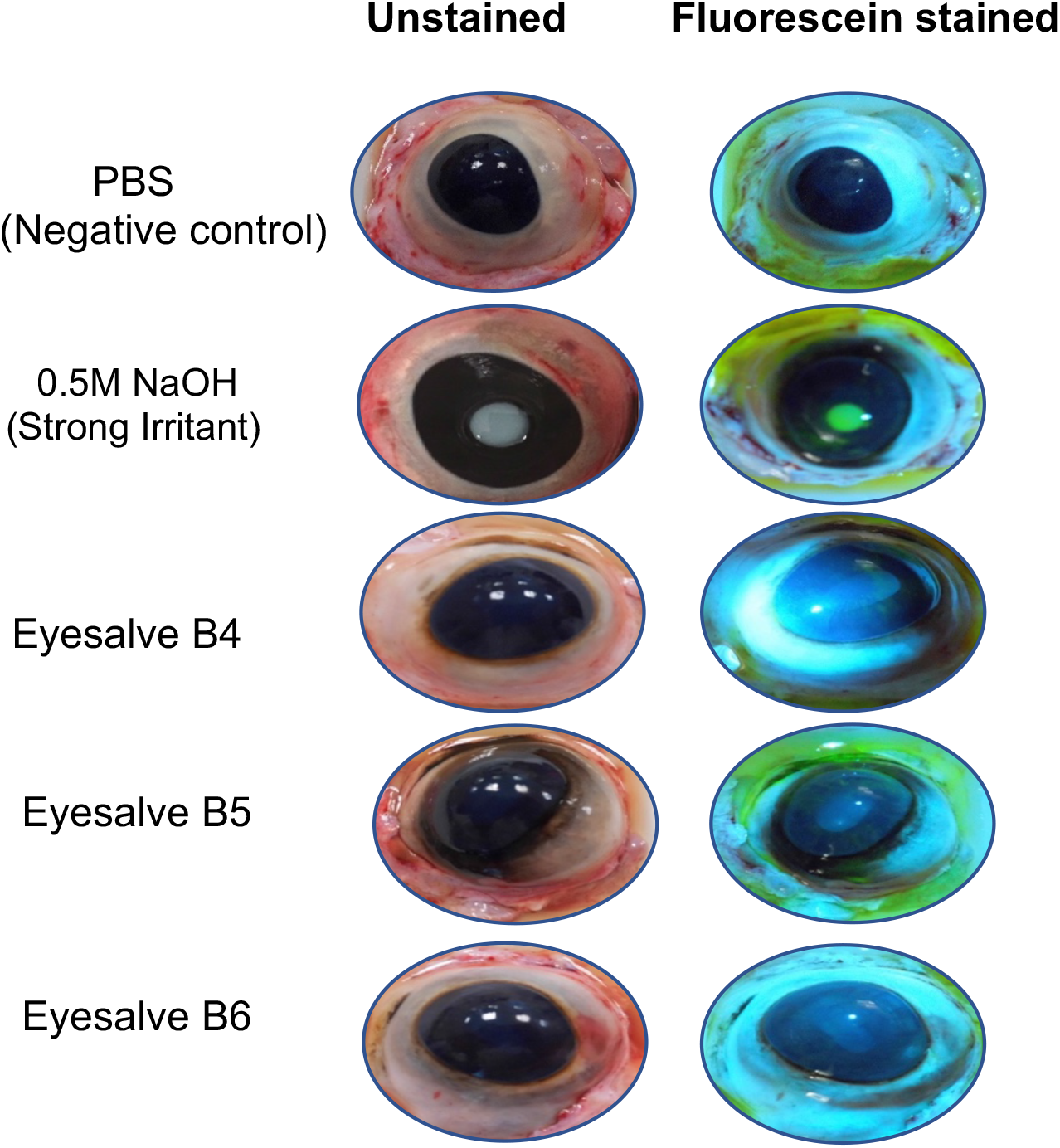
Representative images of bovine eyes treated with different batches of eyesalve. BCOP assay showing unstained and fluorescein stained eye images that were treated with PBS (negative control), 0.5M NaOH (positive control, strong irritant) and different eyesalve batches (B4, B5 & B6).

**Table 1:** Bovine corneal opacity and permeability assay cumulative scores for bovine eyes treated with Bald’s eyesalve and the controls. Six batches of eyesalve (B4, B5, B6, B12, B13, B14) were used.

| Treatment | Mean<br>Opacity<br>score | Mean epithelial<br>integrity score | Cumulative<br>score | Description |
| --- | --- | --- | --- | --- |
| 0.5M NaOH<br>(positive control) | 4 | 1.5 | 5.5 | Severe |
| PBS<br>(negative control) | 0 | 0 | 0 | None |
| B4 | 0 | 0 | 0 | None |
| B5 | 0 | 0 | 0 | None |
| B6 | 0 | 0 | 0 | None |
| B12 | 0 | 0 | 0 | None |
| B13 | 0 | 0 | 0 | None |
| B14 | 0 | 0 | 0 | None |

### Bald’s eyesalve causes slight irritation in a slug mucosal assay

To further determine the safety of the eyesalve, the slug mucosal irritation assay was performed with slugs treated with eyesalve and the weights of the mucus secreted by the slugs before and after treatment were compared (Figure 5A).

**Figure 5:**
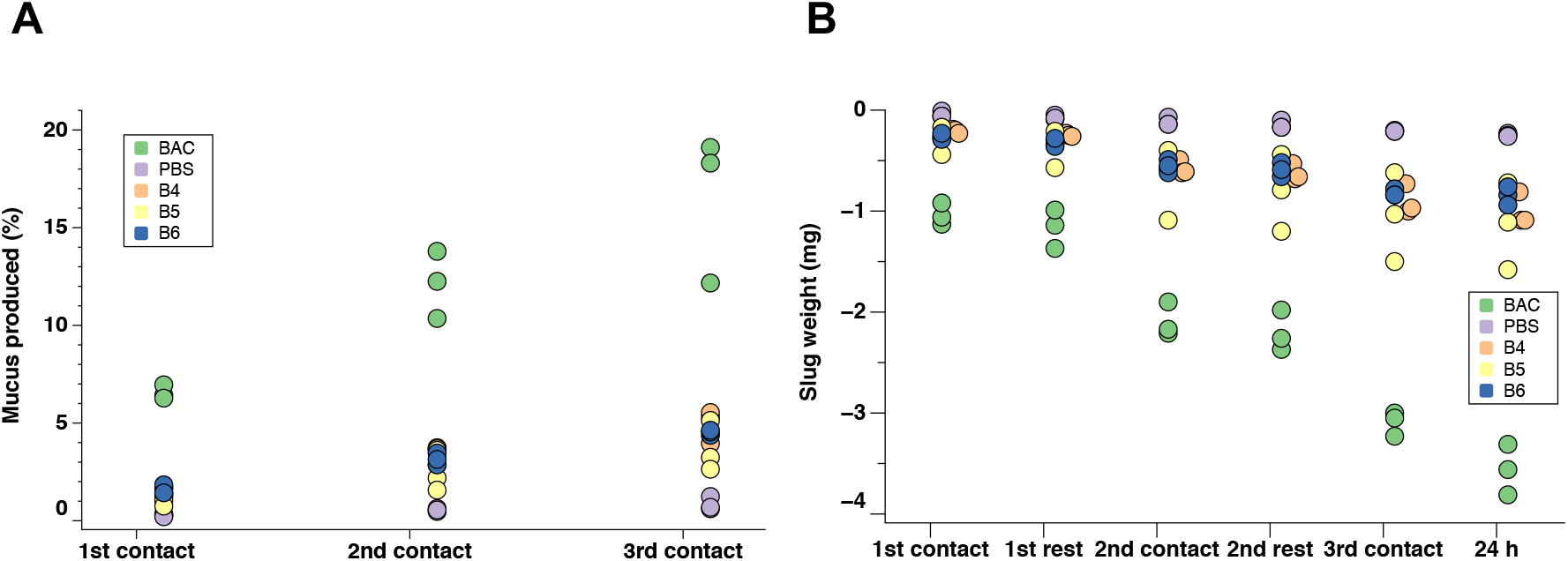
Slug mucosal irritation assay of slugs treated with three batches of eyesalve. A) The percent of mucus produced as a result of irritation after three consecutive 15 minutes contacts with the eyesalve and a rest of 60 minutes between the different contacts. The strong irritant positive control, benzalkonium chloride (BAC) and non-irritant negative control PBS. B4, B5, and B6 are the different batches of the eyesalve. A significant amount of mucus was released in the positive control compared to the eyesalve treated slugs followed by Dunnett’s test for multiple comparison, ANOVA, F_4,10_ = 50.91, p < 0.001, n = 3 replicates) and B) Slug weight loss at the different contacts and after 24h. More weight loss occurred in the positive control compared to the eyesalve treated slugs followed by Dunnett’s test for multiple comparison ANOVA F_4,10_ = 86.63, p < 0.001, n = 3 replicates).

The positive control, benzalkonium chloride, caused severe irritation to the slugs by the third contact period and differed from the eyesalve which caused mild irritation after the third contact period (ANOVA, F_4,10_ = 86.63, p < 0.001). Similarly, the slugs treated with the positive control lost more weight compared to the eyesalve treated slugs (ANOVA, F_4,10_ =50.91, p < 0.001, Figure 5B). Next, the protein concentration of the mucus secreted was quantified using the NanoOrange protein kit. This showed significantly higher amounts of protein in the positive control compared to the eyesalve treated slugs (ANOVA, F_4,30_ = 15.72,, p < 0.002, Figure S2).

### Eyesalve does not interfere with normal wound healing in experimentally wounded mice

Having observed low levels of cytotoxicity and irritation in cell culture, BCOP, and slug mucosal irritation assays, we concluded that an assessment of the safety of the eyesalve in a live vertebrate was justified. We previously reported an approximately 10-fold drop in viable methicillin-resistant *S. aureus* bacteria in tissue biopsies taken from chronic mouse wound infections treated for four hours with the eyesalve (10). In this previous study, we did not treat live mice with the eyesalve or assess any tissue effects, so we aimed to investigate whether the eyesalve would interrupt normal wound healing. We thus used 19 mice, administered a full-thickness surgical excision wound (25) and treated each mouse by applying 100 μl eyesalve or 100 μl sterile water to the wound. The area of the wound size reduced daily by the different treatments (Figure 6A). There was no significant effect of treatment group on wound size at day zero, as wounds were not consistently bigger or smaller in mice assigned to the different treatments (ANOVA, F_3,15_ =1.03, p = 0.41). All wounds were healed to comparable sizes by day 15 irrespective of treatments (ANOVA, F_3,15_ =0.825, p = 0.5). There were no visible signs of irritation or inflammation, thus no reddening of tissue and the mice did not display signs of irritation.

**Figure 6:**
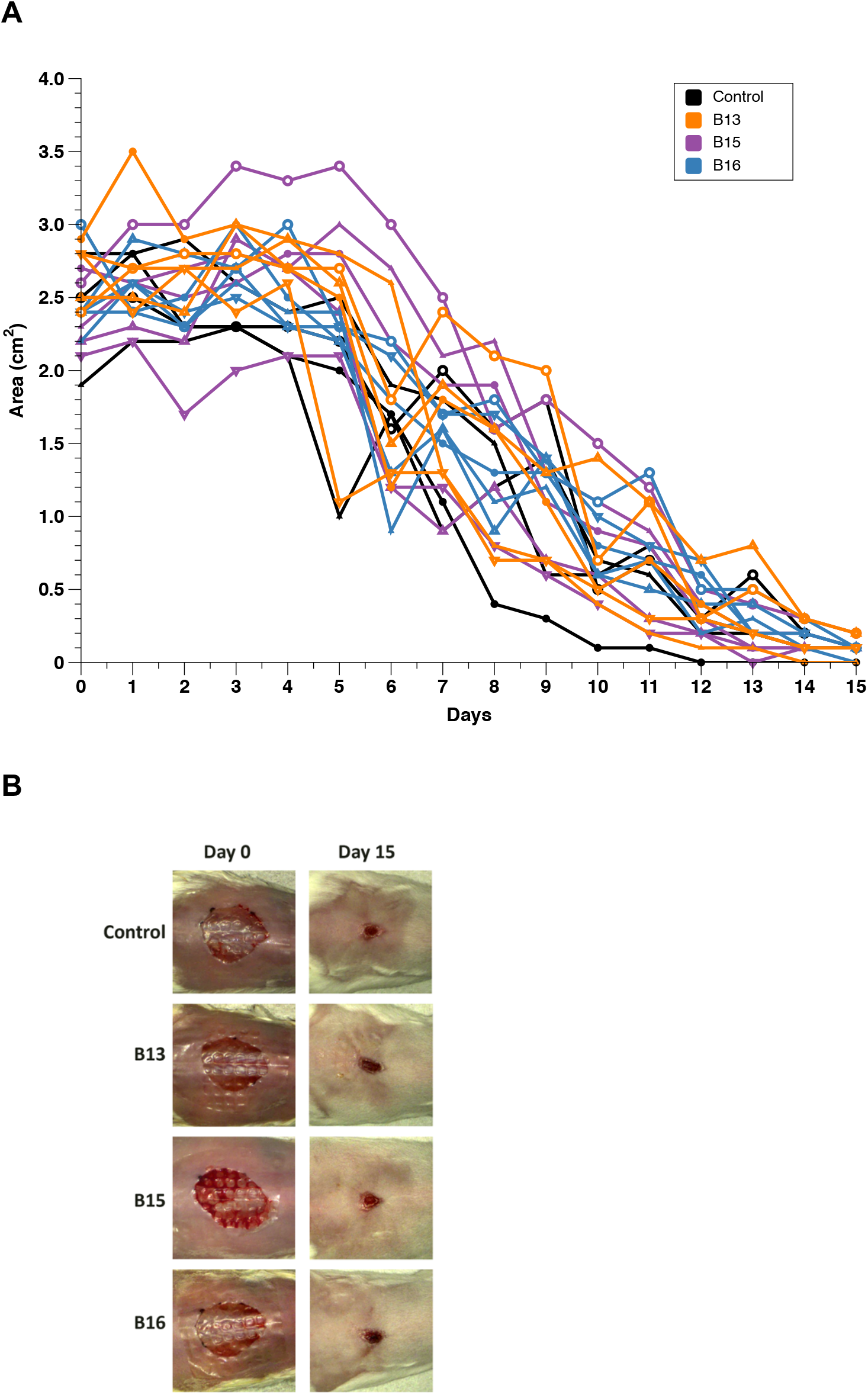
Mice wound closure experiment to determine the safety of Bald’s eyesalve in an *in vivo* model. A) Area of individual mice wounds treated with 100 μl of three batches of eyesalve (B13, B15 and B16) daily for 15 days. The control is sterile water. No difference between wound size by treatment group at day zero (ANOVA, F_3,15_=1.03, p = 0.41) or by day 15 (ANOVA, F_3,15_=0.825, p = 0.5). B) Representative images of the mouse wounds at day 0 and day 15 after treatment with three batches of eyesalve showing a closure of the wounds.

Example of photographs of wounds treated with the eyesalve at days 0 and 15 are shown in Figure 6B. More images of the individual mice treated with eyesalve and the control at different days are presented in Figure S3. We used linear mixed models to compare the rate of wound healing among the four treatments, including mouse identity as a random factor to fit individual relationships between wound area and time for each mouse. There was no difference in healing rates between treatments (testing for an effect of including a time*treatment interaction in the model to allow the slope of the area on time to vary between treatments (*X^2^* = 3.012, p = 0.39, df = 3). Consistent with the analysis of day 0 data, the intercepts of the fitted relationships did not vary between treatments (*X*^2^ = 2.56, p = 0.47, df = 3). However, there was a significant effect of time on wound area, i.e. wounds heal over time (*X*^2^ = 592, p < 0.001, df = 1).

## Discussion

Plant-based remedies have been used in ancient times before the advent of traditional antibiotics and could serve as alternatives in the search for new drugs (6, 9). However, it is necessary that any potential product to be used for the treatment of humans is tested for safety and efficacy.

Previously, we have shown that a medieval remedy, Bald’s eyesalve, possesses antibacterial activity against *S. aureus* in both planktonic cultures, biofilms, and in chronic mouse wounds infected with a methicillin-resistant *S. aureus* strain (10). Recently, we have extended this research by demonstrating the bioactivity of the eyesalve against a range of Gram-positive and Gram-negative organisms in planktonic cultures and biofilms (12).

Here, we show that Bald’s eyesalve is also effective against a multidrug-resistant

*N. gonorrhoeae* strain. *N. gonorrhoeae* is a common cause of ophthalmia neonatorum or neonatal conjunctivitis acquired during delivery from an infected mother. If left untreated, it can lead to corneal perforation and blindness, with neonatal conjunctivitis being a major cause of childhood corneal blindness in developing countries (26–28). With the increasing occurrence of multidrug-resistant strains of bacteria, an alternative treatment for *N. gonorrhoeae* is very valuable and these experiments show that Bald’s eyesalve has the potential to be developed into a viable alternative.

As the safety profile of the eyesalve has not previously been reported, we used a range of models to assess the irritation potential of the eyesalve with the aim of reducing the number of animals used. First, we performed cytotoxic profiling for cell viability with alamarBlue™ using HaCaT and THP-1 cells. The undiluted form of the eyesalve was toxic to the HaCaT cells, as was the positive control, Optrex™ chloramphenicol. This suggests that while the eyesalve can cause damage in the cell lines used, it may not cause significant damage to intact eyes since the undiluted control (chloramphenicol eye drops, 5 mg/mL) caused a similar level of cell death in the skin cells and are accepted for use to treat eye infections. Similar results have been shown in other studies of undiluted plant-based remedies (29, 30). Low numbers of viable cells were observed in THP-1 cells treated with both the undiluted and the 1/10 dilution of Bald’s eyesalve when compared to the chloramphenicol treated controls.

Levels of lactate dehydrogenase, a cytosolic enzyme released when the plasma membrane is damaged, were low in the treated HaCaT and THP-1 cells. However, the levels were much higher in the 1/10 chloramphenicol treated cells than in the 1/10 eyesalve treated cells. It is pertinent to note that cell lines are *in vitro* tools for preliminary screening and do not have the complexity of human tissue (31). Therefore, the results of the alamarBlue™ and LDH assays demonstrate that while the eyesalve causes some damage to cell lines, this may occur at a sufficiently low level to be insignificant *in vivo*.

We thus employed whole-organ and whole-animal models that are used in the consumer healthcare industry to assess the irritation potential of drugs and cosmetic products. The BCOP test, an alternative to the *in vivo* Draize rabbit test, is an *ex vivo* assay that has been used to determine the safety potential of chemicals causing serious eye damage (17, 18). In this assay, the eyesalve did not cause irritation based on the scoring criteria, although some transient mild opacity was observed during the assay, which resolved quickly. Similarly, the slug mucosal irritation assay demonstrated that the eyesalve was only mildly irritating to slugs, which closely mimic irritation of the human mucosa (Figure 5A). Both assays provide evidence of the potential safety of the eyesalve and this has broad implications; while further tests are needed, it seems possible that the eyesalve could potentially be employed in the topical treatment of eye infections and other mucosal site infections. This could be particularly relevant for the therapy of neonatal conjunctivitis caused by *N. gonorrhoeae* infections – a condition with a more serious health and economic burden than the styes for which the eyesalve was originally prescribed.

Since the results from the *in vitro* and *ex vivo* assays were promising, we then tested the safety of the eyesalve in a mouse wound model and showed there was no inhibition of wound closure or signs of irritation. This is important as wound healing poses a serious problem, especially in individuals with chronic non-healing diabetic foot infections (32). As such, being able to design alternative therapies with antimicrobial properties with no effect on wound healing/irritation would be advantageous. This result increases the confidence with which we can suggest Bald’s eyesalve as a candidate for *in vivo* testing of antibacterial potential.

With the increase in antimicrobial resistance, alternatives to antibiotics are urgently needed and Bald’s eyesalve could potentially be used to treat bacterial infections through topical application. This would ideally involve the formulation of a defined natural product cocktail produced under good manufacturing processes to ensure reproducible efficacy and safety. Key candidate for future testing could include diabetic foot ulcers with a treatment cost average of £7800 per infection annually in the UK (33). Other potential treatment target include neonatal conjunctivitis, a major cause of childhood corneal blindness in developing countries (26). Our results are very promising, and the next steps would be to perform rigorous topical testing on healthy humans to determine any potential irritation or toxicity.

## Materials and Methods

### Preparation of Bald’s eyesalve

Bald’s eyesalve was prepared using a standard method as previously reported with garlic, onions, bovine bile and white wine (10). Briefly, the garlic and onions were chopped, pounded together in a pestle and combined with equal volumes of wine (Pennard’s organic dry white, 11% ABV, Avalon Vineyard, Shepton Mallet) and bovine bile salts (Sigma Aldrich). The bovine bile salts were made up to 89 mg/mL in water and sterilised by exposing to UV radiation for ten minutes. The mixture was stored in the dark at 4°C for nine days, after which it was strained, centrifuged and stored in sterilised glass bottles in the dark at 4°C till when used. Several batches of the eyesalve, made on different days, were used for the experiments.

### Media preparation

Gonococcal (GC) broth was prepared by dissolving 3.75 g of protease peptone (Sigma-Aldrich), 0.25 g of potassium phosphate monobasic (Sigma-Aldrich), 1 g of potassium phosphate dibasic (Sigma-Aldrich) and 1.25 g of sodium chloride (Sigma-Aldrich) into 250 mL of distilled water. The GC broth medium was sterilised by autoclaving at 121°C for 15 minutes, cooled to 50°C before adding 250 μl of the iron and 2.5 mL of the glucose supplement aseptically. GC agar was prepared by dissolving 9 g of GC agar base (CM0367, Oxoid) in 250 mL of distilled water, sterilised by autoclaving, and cooled before adding 250 μl of the iron and 2.5 mL of the glucose supplement.

### Disk diffusion assay

*Neisseria gonorrhoeae* NCCP11945 strain was transferred into the GC broth using a sterile loop. 50 μl of the GC broth containing the NCCP11945 strain was transferred to each GC agar plate and was spread evenly using a sterile plastic spreader. For each plate, four sterile 6 mm blank disks were used and 10 μl of each batch of eyesalve was transferred to the blank disk and distilled water was used as a negative control. The plates were incubated in a 5% CO_2_ incubator at 37°C and the zones of inhibition were measured with a ruler after 24 hours and the experiment was performed in triplicate.

### Log reduction assay

The log reduction method was performed according to Bergsson and colleagues (34) with minor modifications as in Churchward *et al*. (2017). *N. gonorrhoeae* strain NCCP11945 was inoculated into the GC broth to an optical density of 0.25-0.30 (520 nm), corresponding to approximately 10^7^ cells per mL. The suspended bacteria (13 μl) was added to 10 μl of each eyesalve and mixed for two minutes. Phosphate buffered saline (PBS) was used as the negative control. After two minutes, 10 μl was transferred to 90 μl of GC broth and serial dilutions performed. The dilutions were plated on a GC agar plate and incubated in 5% CO_2_ at 37°C for 48 hours before counting the colonies.

### Cell culture, media and conditions

Human keratinocytes cell line, HaCaT (P15-P19), a gift from Prof. Mahdad Noursadeghi, University College, London and monocyte-like THP-1 cells (P17-P25), from the European Collection of Authenticated Cell Cultures, ECACC 88081201 were used. HaCaT cells were grown in Dulbecco’s Modified Eagle Medium (DMEM) supplemented with 10% heat inactivated foetal bovine serum (FBS, Labtech, UK), and 1% penicillin-streptomycin (10,000 units/mL penicillin, 10 mg/mL streptomycin, Sigma-Aldrich, UK). THP-1 cells were grown in Roswell Park Memorial Institute (RPMI) medium supplemented with 10% heat inactivated FBS, 1% penicillin-streptomycin, 2 mM glutamine (Sigma-Aldrich, UK). All cell lines were maintained in 5% CO_2_ in a humidified incubator at 37°C with a media change every two days.

### AlamarBlue™ cell viability assay

HaCaT cells (100 μl of 2 x 10^5^ cells/mL) and THP-1 cells (100 μl of 1 x 10^6^ cells/mL) were seeded in 96 well tissue culture plate (Nunc™, Thermofisher) for 24 hours using DMEM and RPMI media respectively supplemented with 10% heat inactivated FBS in 5% CO_2_ incubator at 37°C. The media was aspirated from the seeded HaCaT cells and 100 μl of each eyesalve, Neosporin^®^ antibiotic ointment (neomycin/polymyxin B/bacitracin, 3.5 mg/mL) and Optrex™ chloramphenicol (5 mg/mL) were added to the respective wells. The cell only control media was replaced with 100 μl of DMEM media supplemented with 10% heat inactivated FBS and a media only control was included with the various treatments and left to incubate for 24 hours. For the 1 in 10 dilution, the eyesalve and controls were diluted in DMEM media supplemented with 10% heat inactivated FBS and 100 μl added to each well. As the THP-1 cells are suspension cells, the media (50 μl) was carefully removed with a pipette and replaced with 50 μl of the various treatments. For the diluted treatment, 10 μl was removed and replaced accordingly with the eyesalve and controls. The alamarBlue™ assay was performed according to the manufacturer’s instructions with 10 μl alamarBlue™ (Thermofisher, UK) added to all wells. The absorbance readings at 570 nm and 600 nm after 24 hours of treatment were performed on Tecan SPARK 10M. Three to four replicates per treatment were performed for each batch of eyesalve and controls. The resulting readings were subtracted from the media only controls and converted to percent viability using the recommended formula (35).

### Lactate dehydrogenase cytotoxicity assay

HaCaT cells (100 μl of 1 x 10^5^ cells/mL) and THP-1 cells (100 μl of 2 x 10^5^ cells/mL) were seeded in 96 well tissue culture plate (Nunc™, Thermofisher) for 24 hours using DMEM and RPMI media respectively supplemented with 5% FBS in 5% CO_2_ incubator at 37°C. The various treatments were added to the cells using the approach as per the alamarBlue™ assay, for 24 hours. Four to five replicates per treatment were performed for each batch of eyesalve and controls. Cyquant™ LDH Cytotoxicity Assay Kit (Thermofisher, UK) was used to determine the lactate dehydrogenase released after the treatment following the manufacturer’s instruction. The absorbance was measured at 490 nm and 680 nm, and LDH activity calculated by subtracting the 680 nm value from the 490 nm value. The resulting readings were subtracted from the media only controls before calculating the percent of LDH released indicating cytotoxicity.

### Bovine corneal opacity and permeability (BCOP) assay

Bovine eyes were collected from a slaughterhouse (ABP, Guildford, Surrey, UK) and transported in sterile saline solution. Before the experiment was carried out, the bovine eyes were examined for epithelium detachment, corneal vascularisation and corneal opacity. The test was performed as previously described (36). Briefly, 100 μl of the eyesalve was added and left for 30 seconds. A strong irritant positive control of 0.5 M sodium hydroxide and a non-irritant negative control of PBS were included in each experiment. Each eye was rinsed with 10 mL of saline solution and incubated for 10 minutes in the water bath. The cornea was examined under a cobalt blue filter (465-490 nm, peak 480 nm) following sodium fluorescein treatment (2% (w/v), pH 7.4). Each eye was scored using the scoring matrix in Supplementary Table 1 (17).

### Slug mucosal irritation assay

This was performed as detailed by Lenoir and colleagues (21). Slugs (*Arion* spp.) were collected from a family house garden (in Ham around Richmond riverside, UK). The slugs were fed carrot, lettuce and cucumber purchased from a local supermarket (Sainsbury’s, Penrhyn Road, Kingston upon Thames, UK). Two days before the start of the experiment, the slugs were isolated, weighed and examined for macroscopic injuries. Each slug was placed in individual boxes with paper towels moistened with PBS (pH 7.4). The slugs were kept at 15°C and the body was wet daily with 30 μl PBS. For the experiment, there are three contact periods of 15 minutes with the eyesalve followed by 60 minutes of rest period. Three different batches of eyesalve (100 μl) were used along with a strong irritant positive control of 1% w/v benzalkonium chloride and a non-irritant negative control of PBS. The amount of mucus produced and the weight of the slugs before and after each of the three contact periods were recorded. The mucus produced was converted to percent of the body weight by dividing the weight of the mucus produced by the body weight of the slug before the start of that contact period. The classification model of Lenoir *et al.* 2011 was used to determine the level of irritation caused by the eyesalve (21). Test substances resulting in ≤ 3% mucus production were classified as causing no discomfort. Those between 3% and 8% were classified as mild discomfort, between 8% and 15% was associated with moderate discomfort and severe discomfort for test substances with mucus production > 15% (21). The protein concentration of the mucus released during the irritation test was analysed using NanoOrange protein kit (Invitrogen™) with infinite M200 pro microplate reader with the excitation wavelength of 485 nm and an emission wavelength of 590 nm. The results were extrapolated using a NanoOrange standard curve obtained using bovine serum albumin.

### Mouse wound closure experiments

This study was carried out in strict accordance with the recommendations in the Guide for the Care and Use of Laboratory Animals of the National Institutes of Health. The protocol was approved by the Institutional Animal Care and Use Committee of Texas Tech University Health Sciences Centre (protocol number 07044). Mice wound closure experiments were performed as previously described (25). Briefly, 19 female Swiss Webster mice were fully anaesthetized with Nembutal (pentobarbital), hair was removed with shaving and Nair (Church & Dwight), and lidocaine was injected subcutaneously at the site of excision prior to manually fully excising approximately 1.5 x 1.5cm area of all skin layers. The open wound was then covered with an OPSITE bandage (Smith and Nephew, UK). Images of non-infected wounds were taken at day 0 prior to treating the wounds with 100 μl of the different batches of eyesalve or water (control) with a 25G tuberculin syringe under the bandage to coat and remain on top of the open wound. Each day, the dressing was gently removed, and a further 100 μl of eyesalve or water was added. Images were taken with a Silhouette wound imaging and documentation system (Aranz Medical) every 24 hours after anaesthetizing mice with isoflurane. This continued for 15 days, by which time all the superficial wounds had healed.

### Statistical analysis

All data were analysed with RStudio version 1.1.447 using linear models followed by ANOVA and Dunnett’s test for multiple comparisons. Except stated otherwise, all the results presented are three independent experiments performed in at least triplicate. Raw data and associated R codes will be made available with the accepted version of this manuscript.

## Acknowledgements

We thank Dr Daniel Padfield for advice on mixed models using R. We thank Cerith Harries and Caroline Stewart of the University of Warwick, School of Life Sciences Media Preparation team for preparing the cell culture media used for this work This work was supported by a Diabetes UK Project Grant to FH (ref. 17/0005690) and Jessica Furner-Pardoe is funded by the MRC Doctoral Training Partnership (grant number MR/N014294/1). Julie Bruce is supported by National Institute for Health Research Capability Funding via University Hospitals Coventry and Warwickshire. The funders had no role in study design, data collection and interpretation, or the decision to submit the work for publication.

## Author contributions

CL and FH conceived this study. BOA, RG, CL, MU, KPR, LASS and FH formulated hypotheses and designed experiments. BOA, VN, JFP, RG, AR and FU performed the experiments. BOA and FH analysed the data. BOA drafted the manuscript. JFP, JB, CL, MU, KPR, LASS and FH contributed to manuscript preparation. All authors reviewed and approved the manuscript.

## Supplementary data

**Supplementary Table 1:** Bovine corneal opacity and permeability assay scoring matrix. (modified from Van Erp & Weterings, 1990). Opacity is scored visually based on what is seen with the white light/unstained and epithelial integrity is scored following fluorescein staining visualised with a cobalt blue filtered light.

| Opacity | Score | Epithelial integrity | Score | Cumulative score | Description |
| --- | --- | --- | --- | --- | --- |
| None | 0 | None | 0 | $\leq 0.5$ | None |
| Slight | 1 | Diffuse and weak | 0.5 | 0.6 - 1.9 | Slight |
| Marked | 2 | Confluent and weak | 1 | 2.0 - 4.0 | Moderate |
| Severe | 3 | Confluent and intense | 1.5 | $> 4$ | Severe |

**Figure S1:**
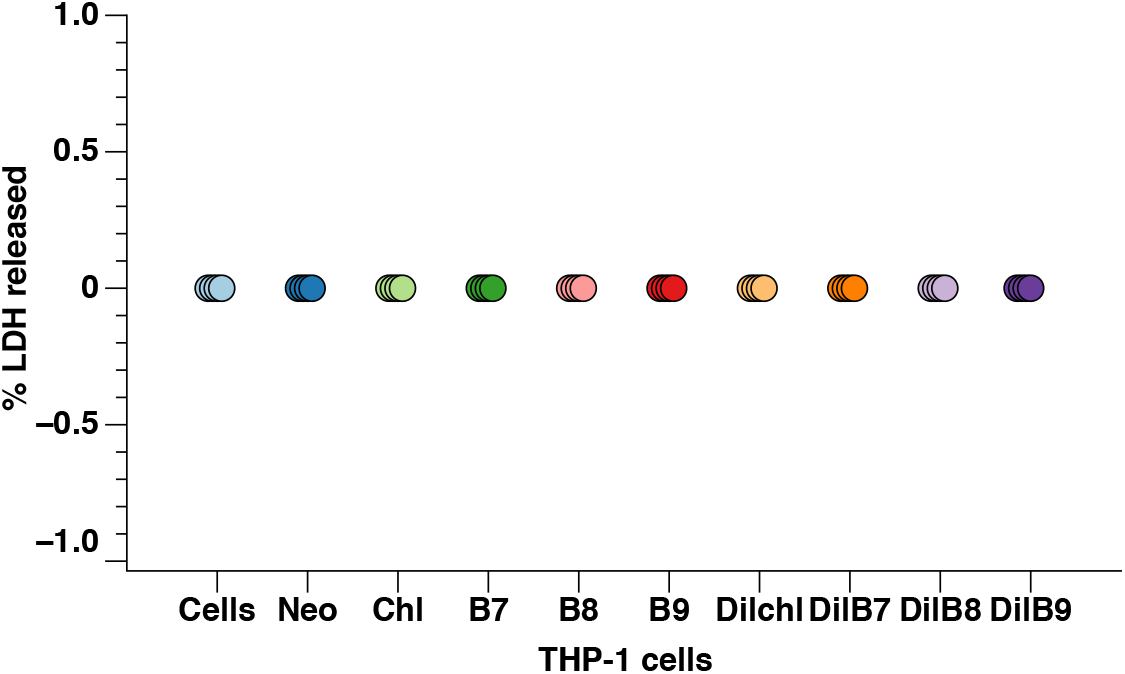
Lactate dehydrogenase assay of THP-1 cells treated with eyesalve. THP-1 cells were treated with three batches of eyesalve (B7, B8 and B9) in the undiluted and diluted (1/10) forms. The controls include cells only (untreated), Neosporin (Neo), a safe antibiotic for wound infections and Optrex™ chloramphenicol (chl) treated cells (n = 4 replicates). The preface “dil” represents cells treated with a 1 in 10 dilution of either the chloramphenicol or the different eyesalve batches.

**Figure S2:**
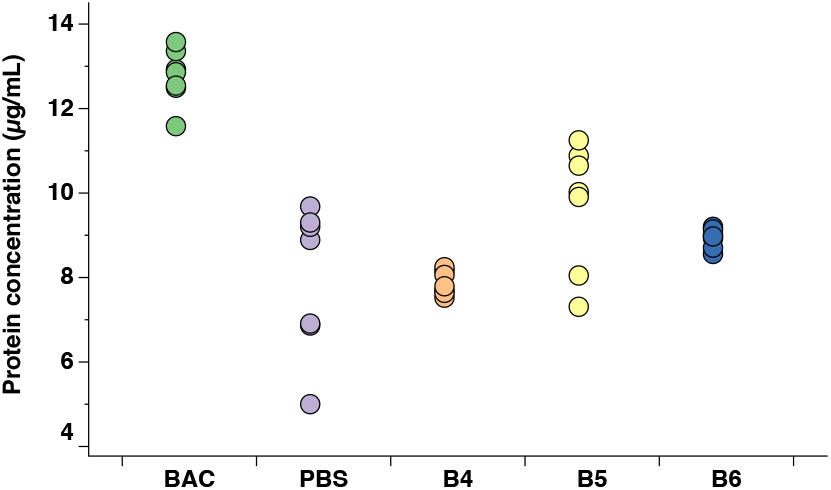
Protein concentration of mucus produced from slugs treated with eyesalve. Slugs were treated with three batches of eyesalve (B4, B5 and B6) and the protein concentration of the mucus measured using the NanoOrange kit. The positive control is benzalkonium chloride (BAC) and the negative control, phosphate buffered saline, PBS. ANOVA found significant higher protein concentration in the positive control compared to the eyesalve treated slugs followed by Dunnett’s test for multiple comparison, F4,30 = 15.72, p < 0.002, n = 7 replicates).

**Figure S3:**
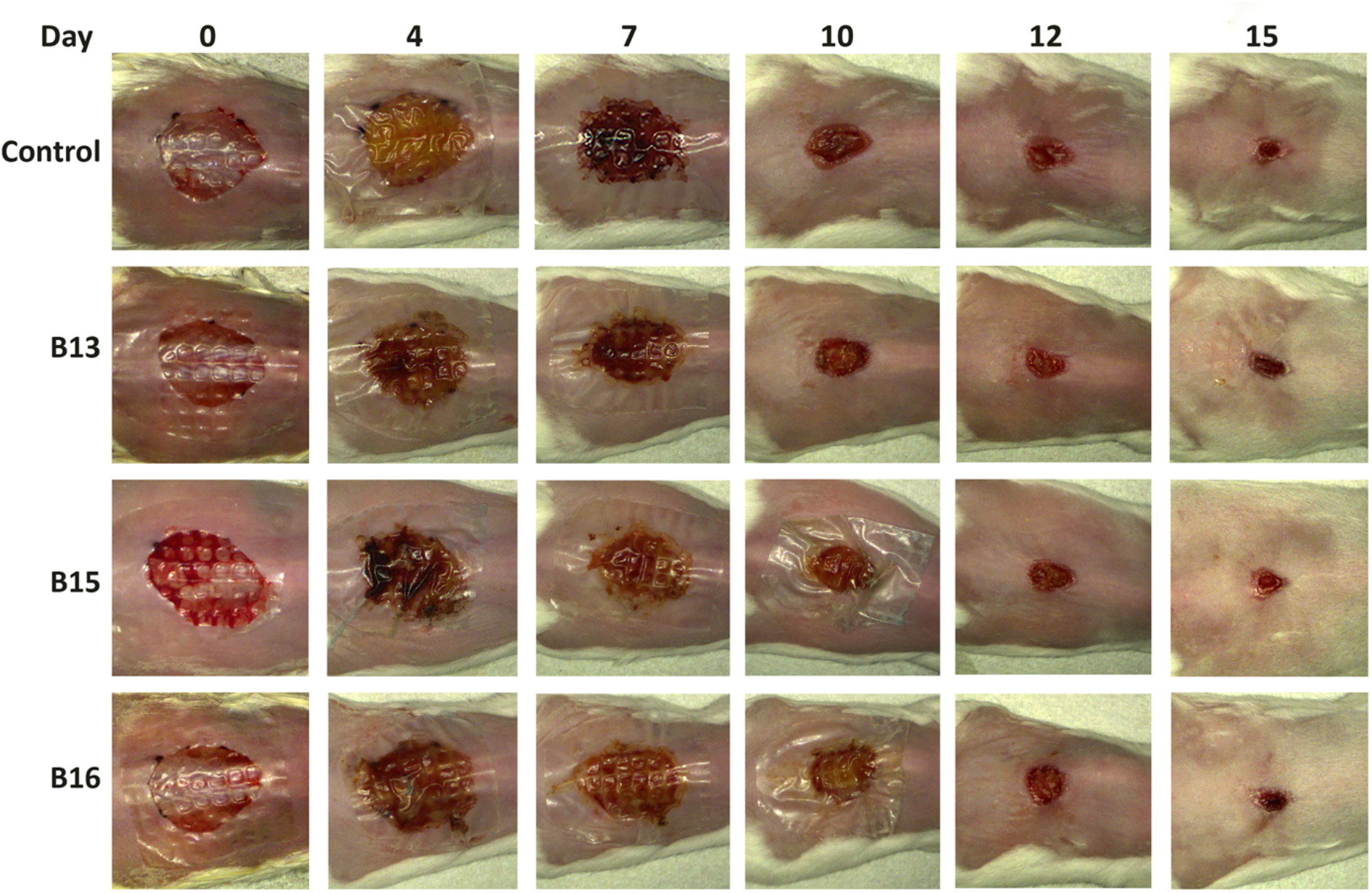
Images of the mouse wounds at different days of treatment with three batches of eyesalve showing closure of the wounds. The control is sterile water.

